## Supplementary material for "Identification of plants functional counterparts of the metazoan Mediator of DNA Damage Checkpoint 1": all supplemental figures

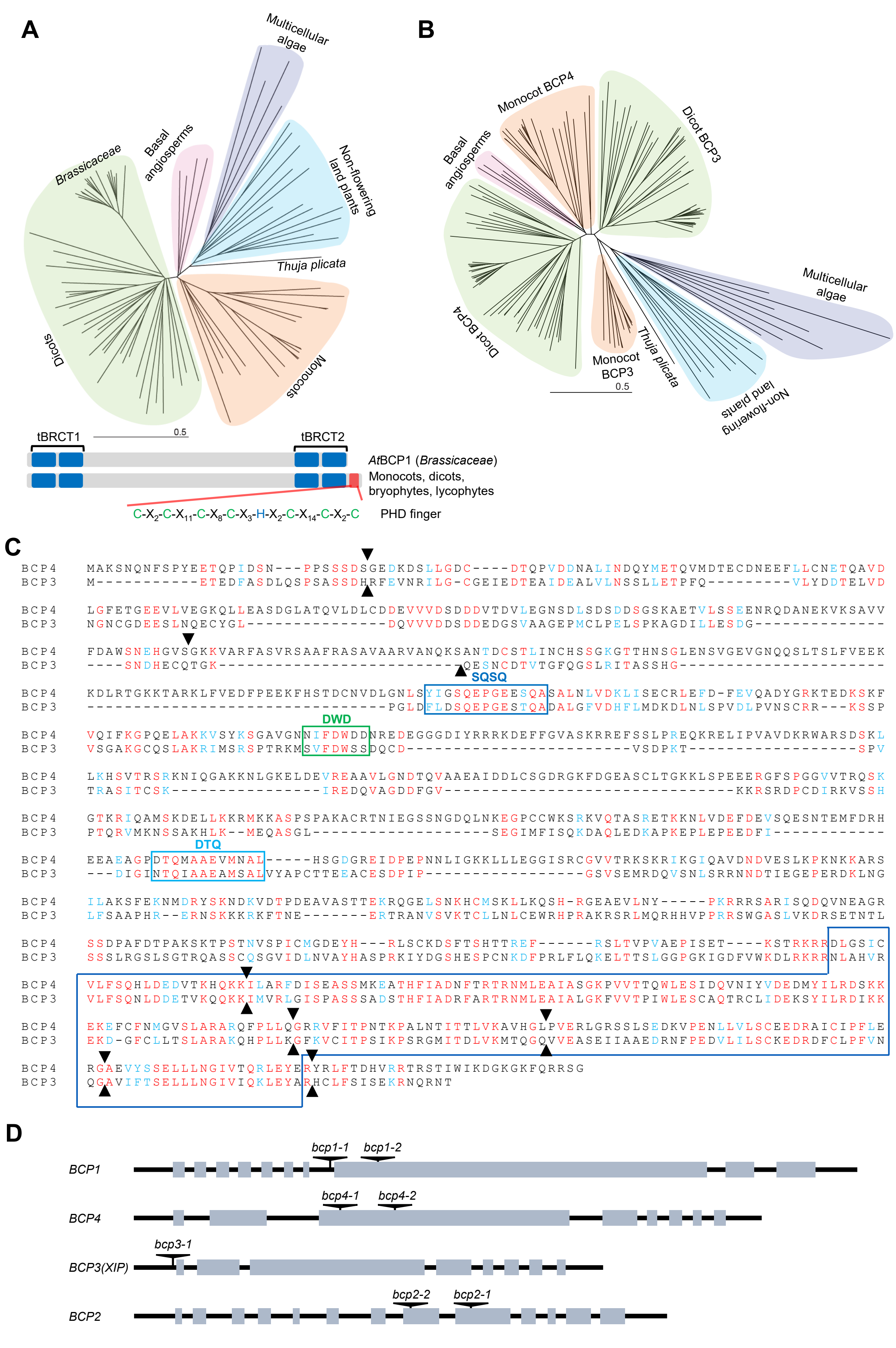


Supporting Figure 1

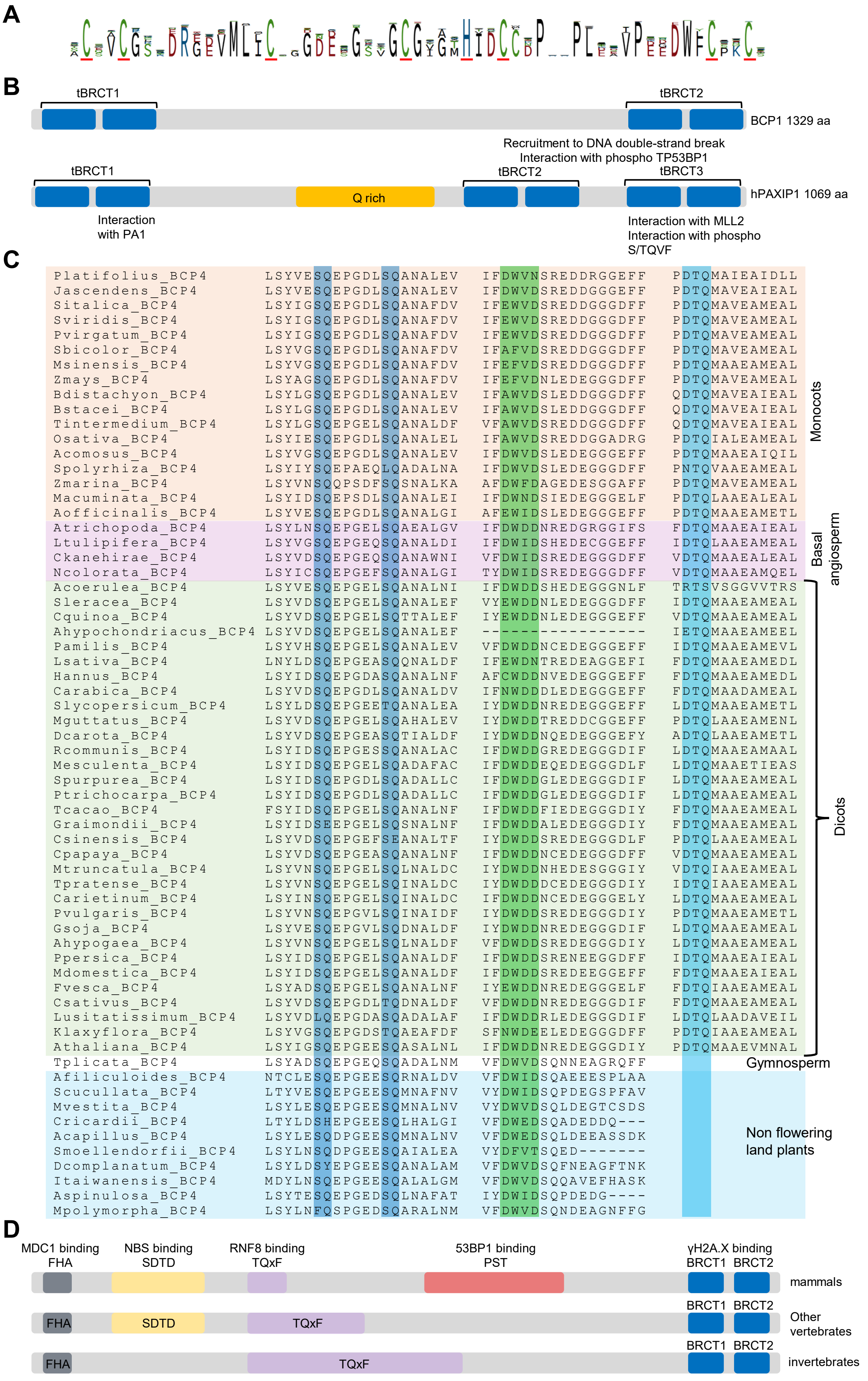


Supporting Figure 2

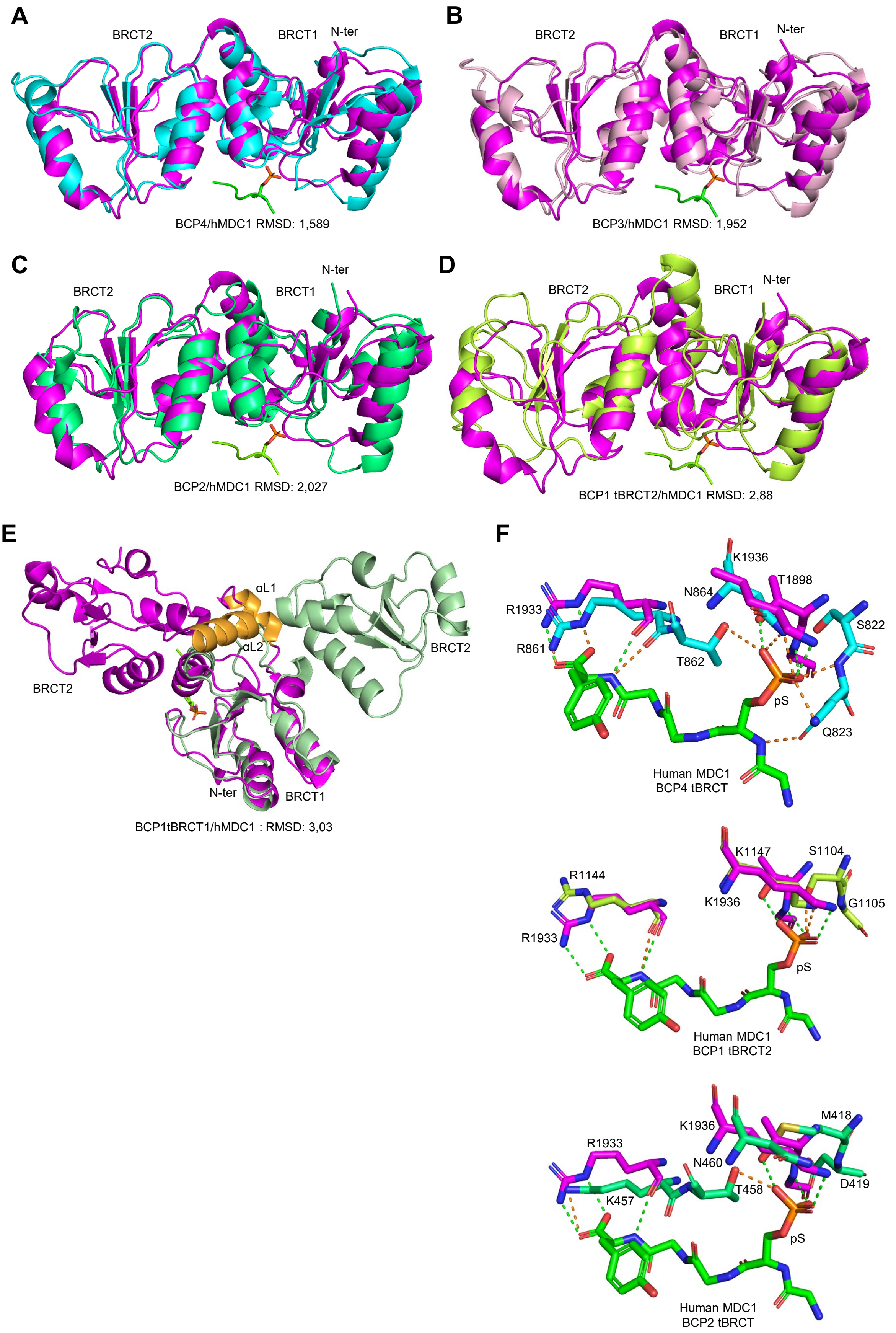


Supporting Figure 3

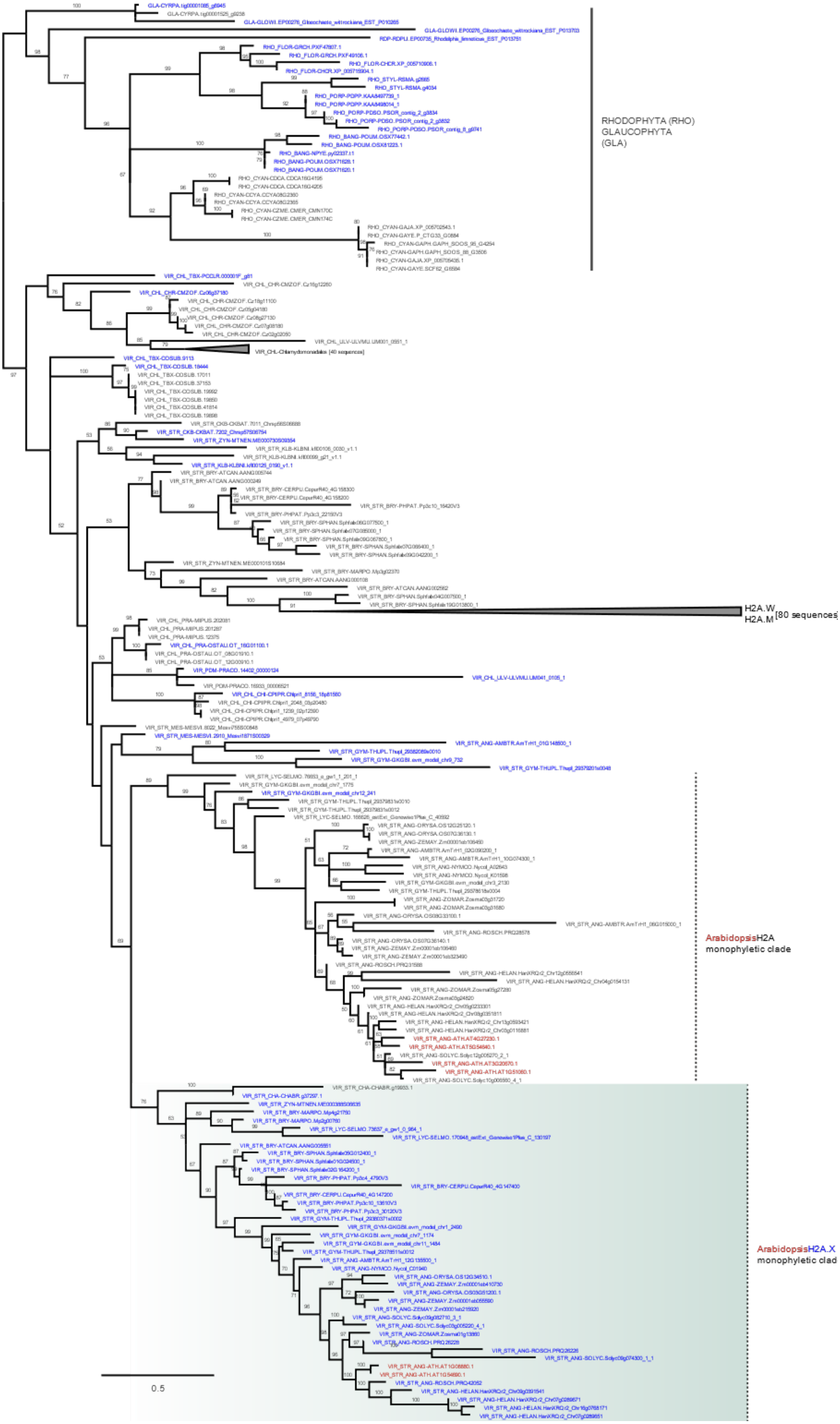


Supporting Figure 4

**A**

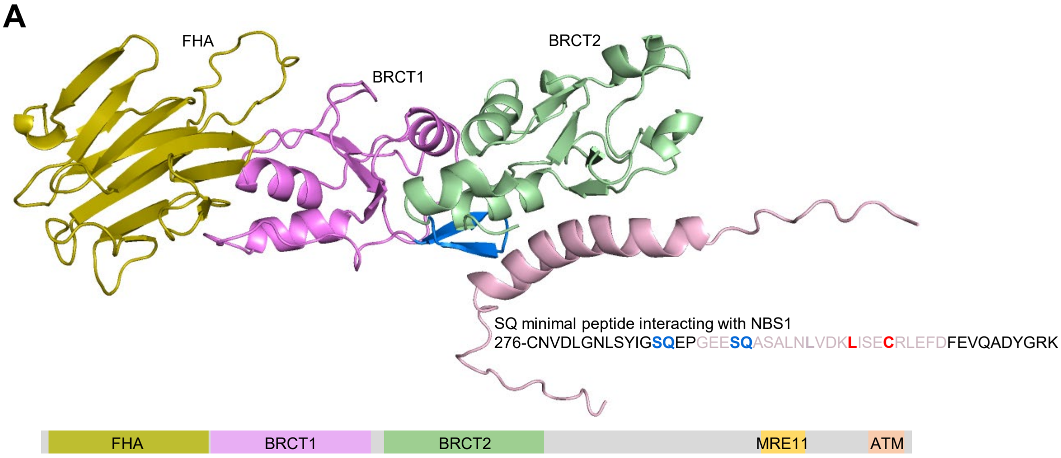


**B**

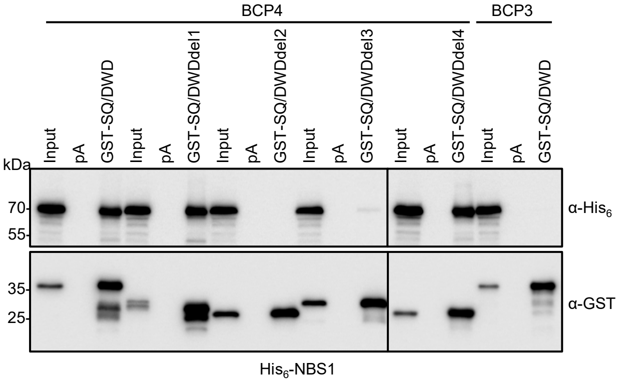


Supporting Figure 5
